# Gene model for the ortholog of *Glyp* in *Drosophila pseudoobscura*

**DOI:** 10.1101/2025.07.18.665514

**Authors:** Megan E. Lawson, Isabel G. Wellik, Viet Le, Evan N. Bennett, Jeffrey S. Thompson, Shallee T. Page, Chinmay P. Rele, Amy T. Hark

## Abstract

Gene model for the ortholog of *Glycogen phosphorylase* (*Glyp*) in the Apr. 2013 (BCM-HGSC Dpse_3.0/DpseGB3) Genome Assembly of *D. pseudoobscura* (GCA_000001765.2). This ortholog was characterized as part of a developing dataset to study the evolution of the Insulin/insulin-like growth factor signaling pathway (IIS) across the genus *Drosophila* using the Genomics Education Partnership gene annotation protocol for Course-based Undergraduate Research Experiences.

## Introduction

*This article reports a predicted gene model generated by undergraduate work using a structured gene model annotation protocol defined by the Genomics Education Partnership (GEP; thegep.org) for Course-based Undergraduate Research Experience (CURE). The following information in quotes may be repeated in other articles submitted by participants using the same GEP CURE protocol for annotating Drosophila species orthologs of Drosophila melanogaster genes in the insulin signaling pathway. (Gruys et al., 2025)*

“In this GEP CURE protocol students use web-based tools to manually annotate genes in non-model *Drosophila* species based on orthology to genes in the well-annotated model organism fruitfly *Drosophila melanogaster*. The GEP uses web-based tools to allow undergraduates to participate in course-based research by generating manual annotations of genes in non-model species (Rele et al., 2023). Computational-based gene predictions in any organism are often improved by careful manual annotation and curation, allowing for more accurate analyses of gene and genome evolution (Mudge and Harrow 2016; Tello-Ruiz et al., 2019). These models of orthologous genes across species, such as the one presented here, then provide a reliable basis for further evolutionary genomic analyses when made available to the scientific community.” (Myers et al., 2024).

We propose a gene model for the *D. pseudoobscura* ortholog of the *D. melanogaster* glycogen phosphorylase (*Glyp*) gene. The genomic region of the ortholog corresponds to the uncharacterized protein XP_001356648.1 (Locus ID LOC4816879) in the Apr. 2013 (BCM-HGSC Dpse_3.0/DpseGB3) Genome Assembly of *D. pseudoobscura* (GCA_000001765.2 - Richards et al. 2005; Drosophila 12 Genomes Consortium 2007). This model is based on RNA-Seq data from *D. pseudoobscura* (SRP006203) and *Glyp* in *D. melanogaster* using FlyBase release FB2023_03 (GCA_000001215.4; Larkin et al., 2021; Gramates et al., 2022; Jenkins et al., 2022).

“The particular gene ortholog described here was characterized as part of a developing dataset to study the evolution of the Insulin/insulin-like growth factor signaling pathway (IIS) across the genus *Drosophila*. The Insulin/insulin-like growth factor signaling pathway (IIS) is a highly conserved signaling pathway in animals and is central to mediating organismal responses to nutrients (Hietakangas and Cohen 2009; Grewal 2009).” (Myers et al., 2024)

“*Glycogen phosphorylase* (*Glyp*; also known as *Glyp, GP, Glp1, DGPH*, FBgn0004507)…has a high degree of amino acid similarity to mammalian enzymes (Tick et al., 1999), and contains repeats that are similar to repeats in several transmembrane proteins including Notch in *D. melanogaster* and *Lin-28* in *Caenorhabditis elegans* (LeMarco et al., 1991). *Glyp* plays a major role in glycolytic activity in skeletal muscle in association with glycogen metabolism and body homeostasis (Yamada et al., 2019). The importance of *Glyp* is evident in that null homozygotes are lethal as full knockouts at the larval stage (Eanes et al., 2006; Yamada et al., 2019). Glyp has also shown to affect development, adult fitness, and aging in *Drosophila* (Bai et al., 2013; Yamada et al., 2019).” (Laskowski et al., 2024).

“*D. pseudoobscura* [NCBI Taxid ID 7237] is part of the *pseudoobscura* species subgroup within the *obscura* species group in the subgenus *Sophophora* of the genus *Drosophila* (Sturtevant 1942; Buzzati-Traverso and Scossiroli 1955). It was first described by Frolova (1929). The *pseudoobscura* species subgroup is endemic to the western hemisphere, where *D. pseudoobscura* is distributed throughout Western North America, Mexico, and Central America (Markow and O’Grady 2005). An additional population of *D. pseudoobscura*, found near Bogota, Colombia, is partially reproductively isolated from the North and Central American populations (Prakash 1972). *D. pseudoobscura* is found primarily in chaparral and temperate forests. *D. pseudoobscura* has been studied extensively in the context of ecological and behavioral genetics, speciation, and genome evolution (Powell 1997).” (Lawson et al., 2024).

### Synteny

The reference gene *Glyp* occurs on chromosome 2L in *D. melanogaster* and is flanked upstream by *CG4259* and *defective proboscis extension response 3* (*dpr3*) and downstream by *tho2* and *CG11723. CG12674* is nested within the second farthest upstream gene, *dpr3*. The *tblastn* search of *D. melanogaster* Glyp-PA (query) against the *D. pseudoobscura* (GenBank Accession: GCA_000001765.2 Genome Assembly database) placed the putative ortholog of *Glyp* within scaffold CH379060 (CH379060.3) at locus LOC4816879 (XP_001356648.1), with an E-value of 0.0 and a percent identity of 88.33%. The putative ortholog is flanked upstream by LOC4817034 (XP_001356649.3) and LOC6902940 (XP_033235331.1), which correspond to *bicoid stability factor* (*bsf*) and *nucampholin* (*ncm*) in *D. melanogaster* (E-value: 0.0 and 0.0; identity: 83.89% and 69.62%, respectively, as determined by *blastp*; Figure 1A; Altschul et al., 1990). The putative ortholog of *Glyp* is flanked downstream by LOC4817125 (XP_001356647.3) and LOC4816777 (XP_001356646.1), which correspond to *tho2* and *CG11723* in *D. melanogaster* (E-value: 0.0 and 4e-91; identity: 81.39% and 47.20%, respectively, as determined by *blastp*). The putative ortholog assignment for *Glyp* in *D. pseudoobscura* is supported by the following evidence: The genes downstream of the *Glyp* ortholog are orthologous to the genes at the same locus in *D. melanogaster* and local synteny is partially conserved, supported by E-values and percent identities, so we conclude that LOC4816879 is the correct ortholog of *Glyp* in *D. pseudoobscura* (Figure 1A).

**Figure 1:**
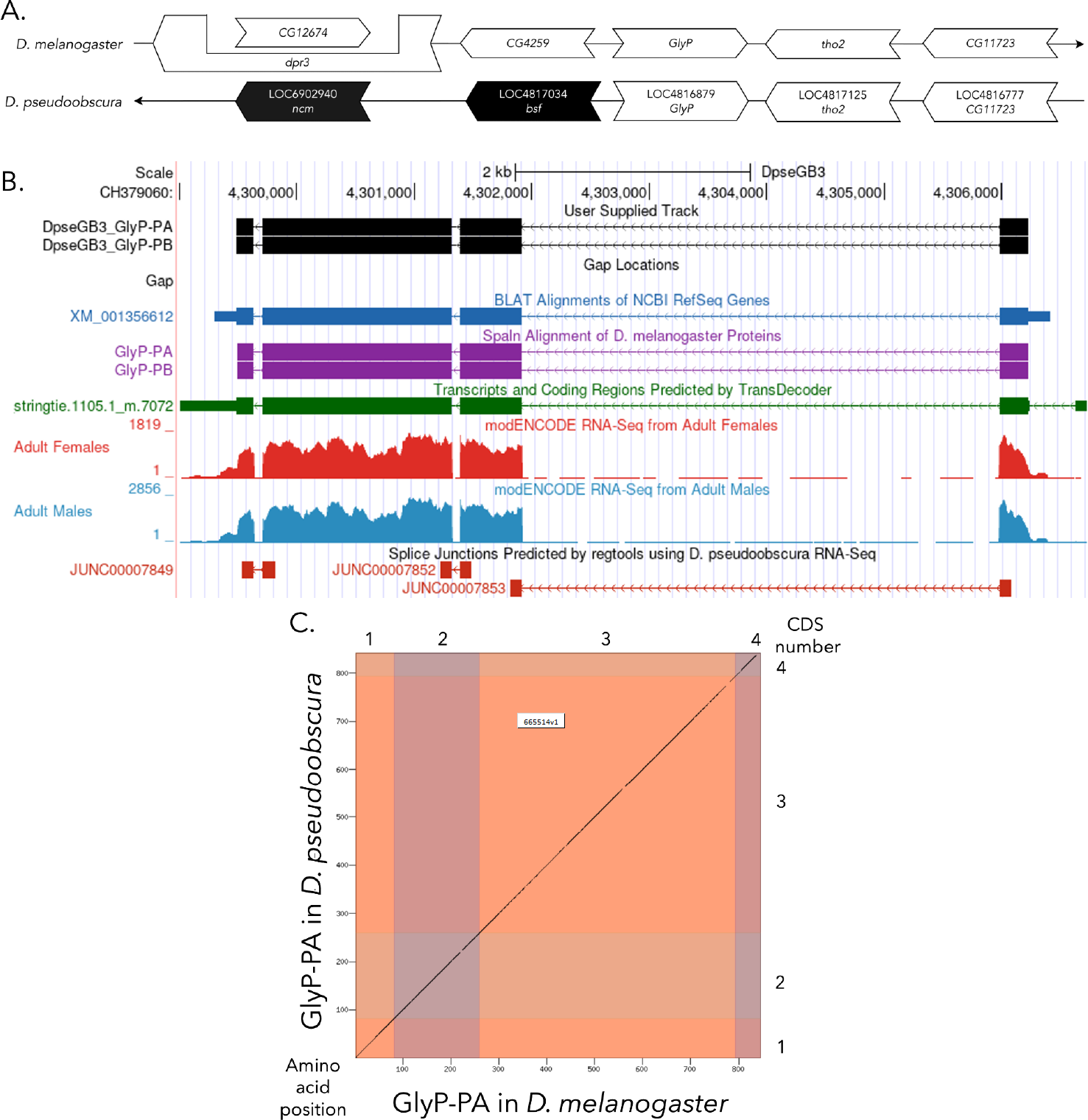
*Glyp* gene model comparison between *Drosophila pseudoobscura* and *Drosophila melanogaster* orthologs (A) Synteny comparison of the genomic neighborhoods for *Glyp* in *Drosophila melanogaster* and *D. pseudoobscura*. Thin underlying arrows indicate the DNA strand within which *Glyp* is located in *D. melanogaster* (top) and *D. pseudoobscura* (bottom). The thin arrow pointing to the right indicates that *Glyp* is on the positive (+) strand in *D. melanogaster*, and the thin arrow pointing to the left indicates that *Glyp* is on the negative (-) strand in *D. pseudoobscura*. The wide gene arrows pointing in the same direction as *Glyp* are on the same strand relative to the thin underlying arrows, while wide gene arrows pointing in the opposite direction of *Glyp* are on the opposite strand relative to the thin underlying arrows. White gene arrows in *D. pseudoobscura* indicate orthology to the corresponding gene in *D. melanogaster*, while black gene arrows indicate non-orthology. Gene symbols given in the *D. pseudoobscura* gene arrows indicate the orthologous gene in *D. melanogaster*, while the locus identifiers are specific to *D. pseudoobscura*. **(B) Gene Model in GEP UCSC Track Data Hub** (Raney et al., 2014). The coding-regions of *Glyp* in *D. pseudoobscura* are displayed in the User Supplied Track (black); coding CDSs are depicted by thick rectangles and introns by thin lines with arrows indicating the direction of transcription. Subsequent evidence tracks include BLAT Alignments of NCBI RefSeq Genes (dark blue, alignment of Ref-Seq genes for *D. pseudoobscura*), Spaln of D. melanogaster Proteins (purple, alignment of Ref-Seq proteins from *D. melanogaster*), Transcripts and Coding Regions Predicted by TransDecoder (dark green), RNA-Seq from Adult Females and Adult Males (red and light blue, respectively; alignment of Illumina RNA-Seq reads from *D. pseudoobscura*), and Splice Junctions Predicted by regtools using *D. pseudoobscura* RNA-Seq (SRP006203). Splice junctions shown have a minimum read-depth of 1000 with >1000 supporting reads indicated in red. **(C) Dot Plot of Glyp-PA in *D. melanogaster* (*x*-axis) vs. the orthologous peptide in *D. pseudoobscura* (*y*-axis).** Amino acid number is indicated along the left and bottom; CDS number is indicated along the top and right, and CDSs are also highlighted with alternating colors. Line breaks in the dot plot indicate mismatching amino acids at the specified location between species.

### Protein Model

*Glyp* in *D. pseudoobscura* has four CDSs within the genome sequence. The unique protein sequence (Glyp-PA) is translated from two mRNA isoforms that differ in their UTRs (*Glyp-RA, Glyp-RB*; Figure 1B). Relative to the ortholog in *D. melanogaster*, the CDS number and protein isoform count are conserved. The sequence of Glyp-PA in *D. pseudoobscura* has 95.84% identity (E-value: 0.0) with the protein-coding isoform Glyp-PA in *D. melanogaster*, as determined by *blastp* (Figure 1C). Coordinates of this curated gene model (*Glyp-RA* and *Glyp-RB*) are stored by NCBI at GenBank/BankIt (accession **BK065278** and **BK065279**, respectively). This gene model can also be seen within the target genome at this TrackHub link:

https://gander.wustl.edu/cgi-bin/hgTracks?db=DpseGB3&lastVirtModeType=default&lastVirtModeExtraState=&virtModeType=default&virtMode=0&nonVirtPosition=&position=CH379060%3A4298987-4306722&hgct_customText=tracktype=bigGenePredvisibility=packbigDataUrl=http://genemodels01.ua.edu/trackhub_hosting/models/DpseGB3/DpseGB3.bb

## Methods

“Detailed methods including algorithms, database versions, and citations for the complete annotation process can be found in Rele et al. (2023). Briefly, students use the GEP instance of the UCSC Genome Browser v.435 (https://gander.wustl.edu; Kent WJ et al., 2002; Navarro Gonzalez et al., 2021) to examine the genomic neighborhood of their reference IIS gene in the *D. melanogaster* genome assembly (Aug. 2014; BDGP Release 6 + ISO1 MT/dm6). Students then retrieve the protein sequence for the *D. melanogaster* reference gene for a given isoform and run it using *tblastn* against their target *Drosophila* species genome assembly on the NCBI BLAST server (https://blast.ncbi.nlm.nih.gov/Blast.cgi; Altschul et al., 1990) to identify potential orthologs. To validate the potential ortholog, students compare the local genomic neighborhood of their potential ortholog with the genomic neighborhood of their reference gene in *D. melanogaster*. This local synteny analysis includes at minimum the two upstream and downstream genes relative to their putative ortholog. They also explore other sets of genomic evidence using multiple alignment tracks in the Genome Browser, including BLAT alignments of RefSeq Genes, Spaln alignment of *D. melanogaster* proteins, multiple gene prediction tracks (e.g., GeMoMa, Geneid, Augustus), and modENCODE RNA-Seq from the target species. Detailed explanation of how these lines of genomic evidenced are leveraged by students in gene model development are described in Rele et al. (2023). Genomic structure information (e.g., CDSs, intron-exon number and boundaries, number of isoforms) for the *D. melanogaster* reference gene is retrieved through the Gene Record Finder (https://gander.wustl.edu/~wilson/dmelgenerecord/index.html; Rele et al., 2023).

Approximate splice sites within the target gene are determined using *tblastn* using the CDSs from the *D. melanogaste*r reference gene. Coordinates of CDSs are then refined by examining aligned modENCODE RNA-Seq data, and by applying paradigms of molecular biology such as identifying canonical splice site sequences and ensuring the maintenance of an open reading frame across hypothesized splice sites. Students then confirm the biological validity of their target gene model using the Gene Model Checker (https://gander.wustl.edu/~wilson/dmelgenerecord/index.html; Rele et al., 2023), which compares the structure and translated sequence from their hypothesized target gene model against the *D. melanogaster* reference gene model. At least two independent models for a gene are generated by students under mentorship of their faculty course instructors. Those models are then reconciled by a third independent researcher mentored by the project leaders to produce the final model. Note: comparison of 5’ and 3’ UTR sequence information is not included in this GEP CURE protocol.” (Gruys, et al. 2025)

## Supplemental Files

1. Zip file containing a FASTA, PEP, GFF files for the gene model
2. Figure 1 in high resolution

### Metadata

Bioinformatics, Genomics, *Drosophila*, Genotype Data, New Finding

## Supporting information

Gene model data files

## Acknowledgements

We would like to thank Wilson Leung for developing and maintaining the technological infrastructure that was used to create this gene model and Laura K. Reed for overseeing the project. Thank you to FlyBase for providing the definitive database for *Drosophila melanogaster* gene models.

